# Application of Machine Learning Tools for Waterbird Colony Monitoring Provides Gains in Precision and Temporal Efficiency

**DOI:** 10.64898/2026.07.01.735369

**Authors:** Anna Vallery, Krish Kabra, R. E. Gibbons, Hank Arnold, Nicholas Minnich, Arko Barman

**Author notes:** Corresponding author, 5902 Lake Washington Blvd. S, Seattle, WA 98118.

## Abstract

Waterbirds serve as important indicators of both aquatic and terrestrial ecosystem health, making effective monitoring essential for tracking population health and identifying potential causes of decline. Drones have provided opportunities to overcome historic waterbird monitoring challenges, but the expertise and time required for manual image analysis creates a major bottleneck. Recent advances in deep learning-based object detection have enabled rapid, automatic detection of features in complex ecological imagery, though applications have largely been limited to single-species colonies, and practitioners lack quantitative comparisons of annotation time and accuracy across different levels of automation. We systematically compared four waterbird monitoring approaches using identical survey areas from Chester Island, a mixed-species colony in Matagorda Bay, Texas, in 2025: (1) traditional ground-based counts, (2) manual drone imagery-based counts, (3) computer-assisted counts using pre-annotations from an object detector with manual human verification (Human+ML), and (4) fully automated counts using object detector annotations (ML-only). We trained a YOLOv10 object detection model on manually annotated imagery of Chester Island in 2021 and applied it to the 2025 imagery. Manual drone annotation detected 6,530 birds in 40.5 hr and served as the primary reference standard. Human+ML detected 5,826 birds (89% of manual) in 7.7 hr, an 81% reduction in annotation time. ML-only detected 5,679 birds (87% of manual) in approximately 46 min, a 98% reduction. Ground counts recorded 5,868 birds (90% of manual). Detection generalized well across species while classification depended heavily on training data and morphological distinctiveness. The Human+ML workflow emerged as a practical middle ground, providing practitioners with empirical data to evaluate partial versus full automation strategies based on monitoring objectives.

**LAY SUMMARY:**

- Conservation programs need accurate counts of nesting waterbirds, but analyzing drone images by hand has become a major bottleneck, slowing the availability of monitoring data for use.
- We compared four ways to count waterbirds at a large nesting colony in coastal Texas where many species nest together: traditional ground-based counts, manual annotation of drone imagery, computer-assisted annotation, and fully automated annotation using a trained object detection model.
- Detection generalized well across species while classification depended on training data availability and morphological distinctiveness.
- Pairing automated detection with site-specific or human classification offers a practical path forward for monitoring mixed-species colonies.

## INTRODUCTION

Effective conservation of wildlife species depends on monitoring programs that can reliably characterize population status and inform management decisions over time. Waterbirds, including seabirds, shorebirds, and wading birds, serve as important indicators of both aquatic and terrestrial ecosystem health due to their sensitivity to environmental changes (Kushlan 1993, Amat and Green 2010, Green and Elmberg 2014). These are also among the most threatened groups of birds, making effective monitoring essential for tracking population health and identifying potential causes of decline (Croxall et al. 2012).

Many waterbird species nest in large, mixed-species colonies, often on nearshore islands with unique breeding bird assemblages dependent on location and habitat structure. This breeding behavior provides opportunities for biologists to study populations that are otherwise challenging to monitor effectively. Methods for surveying these colonies have primarily been traversing the areas on foot, circumnavigating islands via boat, or aerial surveying in small, crewed aircrafts, dependent on species of interest, location, and budget constraints (Green et al. 2010, Ortego et al. 2011, Prosser et al. 2023). Each of these survey methods has associated challenges, including difficulties with accessing colonies, time available for monitoring, and costs associated with the overall monitoring effort (Edney et al. 2023). Potential impacts of monitoring efforts to the birds and habitat of interest are also a concern. Surveying waterbird colonies by foot, for example, can cause temporary adult nest abandonment, which can expose vulnerable chicks to the elements or cause predation of eggs or chicks (Carney and Sydeman 1999). Boat-based surveys often have a low vantage point, which can result in underestimates of breeding populations, particularly on larger and/or higher islands or in areas with taller vegetation (Wetlands International 2010). Researchers often prefer crewed aerial surveys due to the limited disturbance of the colony of interest (Hillman et al. 2015) and excellent vantage points, but this type of survey can be prohibitively expensive and requires access to an airport and favorable weather. Both crewed aerial surveys and boat-based surveys also put biologists at risk - airplane crashes and boating accidents have been found to be the largest causes of mortality and injury among biologists in the field (Sasse 2003).

Uncrewed aerial systems (UAS, drones) have become integral tools in wildlife management over the past decade (Jones et al. 2006, Watts et al. 2010, Nowak et al. 2018), particularly for waterbird monitoring (Chabot et al. 2015, Han et al. 2017). Drones have provided opportunities to overcome those historic waterbird monitoring challenges and have successfully been used to monitor a variety of species and to answer a range of questions about those species (Edney and Wood 2021). The advancement of drone technology and its accessibility allows researchers to remain safely on the ground while operating in and around study areas with less cost and greater ease than traditional methods of aerial surveys. Additionally, drone-based surveys can produce more accurate count estimates than ground-based methods, particularly for ground-nesting species in open habitats (Hodgson et al. 2018, Edney et al. 2023, Arnberg et al. 2026), though the relative accuracy of each method varies with nesting guild and vegetation structure (Prosser et al. 2023).

Nevertheless, while drones have enhanced waterbird monitoring, they have also introduced novel challenges. Specifically, most seabird monitoring studies using drones (92 out of 114 studies) have relied on manual image analysis, where researchers examine images and make counts or measurements themselves (Edney et al. 2023). The expertise and time required for manual image analysis creates a major bottleneck, as localizing and classifying species from hundreds to thousands of aerial images is time-consuming. Even at small, single-species colonies, post-processing of drone imagery can take more than twice as long as the drone surveys themselves (Scarton and Valle 2020). This rate-limiting step has driven interest in both semi-automated and fully-automated methods for UAS image analysis (Edney et al. 2023).

Applying automation to the analysis of aerial imagery has successfully improved monitoring efficiency for various waterbird colony locations and species of interest, including significant reductions in image processing time (Abd-Elrahman et al. 2005, Descamps et al. 2011, Groom et al. 2013, Ratcliffe et al. 2015). More recently, advances in deep learning-based object detection, a sub-field of machine learning (ML) and artificial intelligence (AI), have enabled rapid, automatic detection of features in complex ecological data, including waterbird localization and classification in mixed-species colonies with promising results (Christin et al. 2019, Hong et al. 2019, Jones et al. 2020, Hayes et al. 2021, Weinstein et al. 2022, Kabra et al. 2022, Edney et al. 2023). In one study, for example, what took three weeks to manually analyze (approximately 21,000 birds) was processed by a convolutional neural network (CNN)-based object detector in only 4.5 hr (Kellenberger et al. 2021).

Despite these advances, some key limitations remain. First, applications of object detection for automated waterbird monitoring have been restricted to single-species colonies or in locations with a few, visually distinct species (Chabot and Francis 2016, Hayes et al. 2021). Second, practitioners lack quantitative comparisons of annotation time and accuracy across different levels of automation. Though fully automated methods have demonstrated dramatic time savings, the relative contributions of automated localization (i.e. dotting or boxing) versus automated classification remain unclear. Furthermore, as most studies evaluate either traditional ground surveys or drone-based methods in isolation, there is no direct comparison of accuracy-efficiency trade-offs across the full suite of available approaches. This knowledge gap hinders informed decision-making about incremental adoption of these tools, particularly for conservation programs managing single-species colonies that only need a detector, mixed-species colonies where full automation may not yet be feasible, or those that require maintaining some similarity with historical monitoring protocols.

We address these limitations using colonial waterbird monitoring along the Texas Coast as a model system. Texas coastal colonies present monitoring challenges common to mixed-species waterbird assemblages globally: more than 25 species nest with intermixed ground and shrub nests on nearshore islands, ranging from large, morphologically distinct birds like *Pelecanus occidentalis* (Brown Pelican) to visually similar tern species including *Thalasseus maximus* (Royal Tern) and *Thalasseus sandvicensis* (Sandwich Tern) (Figure 1). This species diversity, combined with varying nesting densities and background substrates, provides an ideal test case for evaluating object detector performance across morphological variation and potential species-level confusion patterns. Additionally, the Texas Colonial Waterbird Survey, one of North America’s longest-running waterbird monitoring programs (Ortego et al. 2011), provides established field protocols and historical context for comparing traditional and emerging monitoring methods.

**Figure 1.**
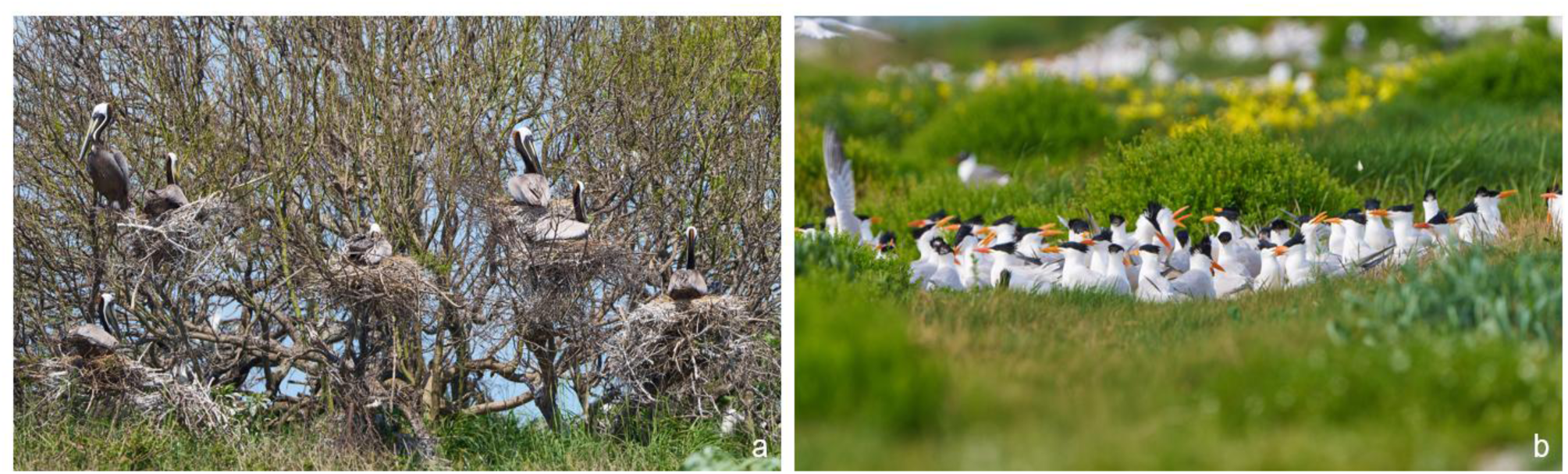
Mixed-species colonial waterbird nesting on Chester Island, Matagorda Bay, Texas. (**A**) *Pelecanus occidentalis* (Brown Pelican) nesting in shrubs. (**B**) Dense *Thalasseus maximus* (Royal Tern) nesting colony on open substrate, representative of ground-nesting species that are readily visible in aerial surveys but challenging to count individually from the ground.

*Alt Text: Two photos taken on Chester Island. On the left is a photo of Brown Pelican adults sitting on large nests of sticks in a barren tree. On the right is a photo of a group of adult Royal Tern sitting together on the ground, surrounded by short, grassy vegetation and slightly taller shrubby vegetation. There are yellow flowers and more birds blurry in the background*.

We compared four waterbird monitoring approaches using identical survey areas from Chester Island, a mixed-species colony in Matagorda Bay, Texas, in 2025: (1) traditional ground-based counts, (2) manual drone imagery-based counts using human-only annotations, (3) computer-assisted drone imagery-based counts using pre-annotations from an object detector with manual human verification, and (4) fully automated drone imagery-based counts using object detector annotations. To enable automated detection of waterbirds in drone imagery, we trained a YOLOv10 object detection model (Wang et al. 2024) on manually annotated imagery of Chester Island in 2021. This model was then applied to the 2025 imagery. For each annotation approach, we obtained both a species-level waterbird count and the time required to complete annotations, enabling direct comparison of workflow output and efficiency. We observed that (1) fully automated methods would substantially reduce processing time relative to manual annotation while maintaining comparable detection rates for most species, (2) semi-automated methods combining automated detection with manual classification would achieve higher classification accuracy than fully automated approaches with meaningful time savings over manual annotation, and (3) classification accuracy under full automation would vary among species as a function of morphological distinctiveness and representation in the training dataset. This study provides practitioners with empirical data to evaluate the relative benefits of partial versus full automation strategies based on their monitoring objectives, available expertise, species assemblage complexity, and requirements for continuity with historical survey methods.

## METHODS

### Study Area

The study site was Chester Island (Figure 2), a 28-hectare dredge-material island that formed in 1962 as a byproduct of the Matagorda Ship Channel creation. Chester Island is located in Matagorda Bay along the Texas mid-coast (28.4167°N, 96.4000°W). It is one of the largest and most productive colonial waterbird nesting islands in the Gulf of Mexico. Managed by the National Audubon Society, Chester Island has been surveyed annually as part of the Texas Colonial Waterbird Survey, with breeding bird data available since the 1970s. Approximately 25 waterbird species have been documented nesting on Chester Island, including Brown Pelican, Laughing Gull, several tern and heron species, and other wading birds.

**Figure 2.**
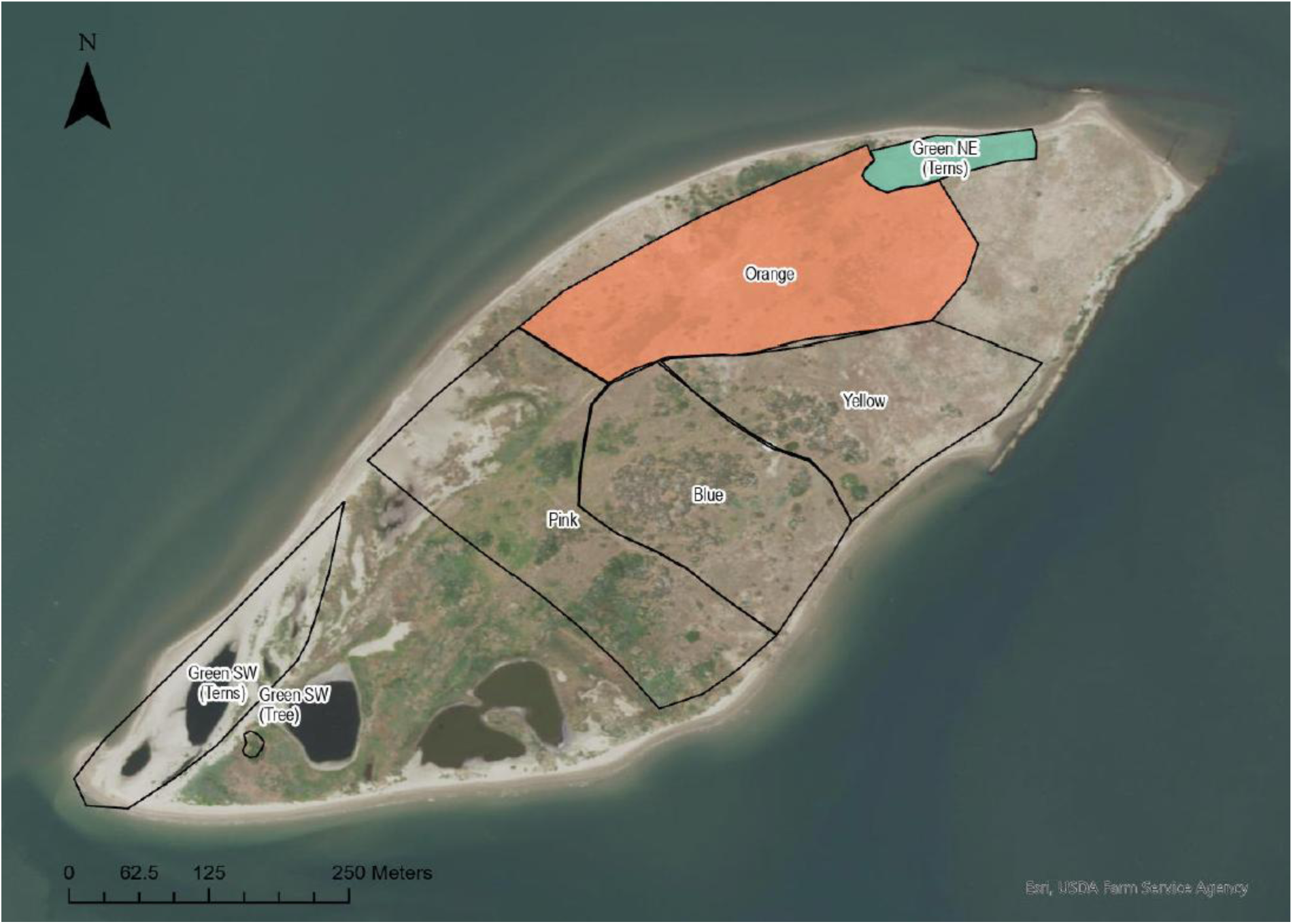
Survey zones on Chester Island, a 28-ha colonial waterbird nesting island in Matagorda Bay, Texas (28.4167°N, 96.4000°W). Outlined polygons delineate the areas used by ground-based teams during the annual Texas Colonial Waterbird Survey census. For this study, comparisons among ground-based, manual drone, Human+ML, and ML-only monitoring methods focused on two zones: the Orange polygon (Brown Pelican, Laughing Gull, and tree-nesting wading birds) and the Green NE polygon (terns, with some additional Brown Pelican and Laughing Gull). Imagery: Esri, USDA Farm Service Agency.

*Alt Text: A map of Chester Island using satellite imagery with black polygons demonstrating the survey areas that ground teams use. Two of the polygons towards the NE side of the island are shaded with color, one larger polygon in orange and another in green*.

Specific survey areas within Chester Island were selected based on previously established count areas used during annual ground-based field surveys (Figure 2). Two of these areas, designated “Orange” and “Green NE,” were used for direct method comparisons. The orange area included nesting habitat for Brown Pelican, Laughing Gull, and tree-nesting wading birds, while the green area was primarily nesting habitat for terns, with some additional Brown Pelican and Laughing Gull individuals present. These two survey areas were selected as they included individuals from each nesting species and demonstrated the variety of habitat types on Chester Island.

## Field Methods

### Ground-based surveys

Chester Island is accessed by boat. During an established time frame set during the peak of nesting abundance, observer teams walk through the nesting colony and count and estimate nesting pairs within pre-designated areas using visual detection. For ground-nesting species such as Royal and Sandwich terns, a 10-ft ladder is brought to the edge of the nesting group to provide the observer with a higher viewing angle of the tightly packed nests.

### Drone Surveys

The island was also surveyed using a DJI Matrice 300 RTK quadcopter with a Zenmuse P1 46 megapixel camera (DJI, Shenzhen, China) during each Texas Colonial Waterbird Survey period. The drone flew at 30-40 m altitude, capturing photos every 0.7 s. The camera was positioned at 90° nadir (straight down) with flight speeds typically 25 km per hr to avoid obvious disturbance signs and ensure clear imagery. Transects covered the entire island with 35% overlap between adjacent transects, ensuring complete coverage and capturing each bird in at least one clear image. All images were georeferenced to enable direct spatial comparisons between methods. Survey imagery was processed into orthomosaic images, then rendered into 10,000 × 10,000-pixel images.

### Object Detector Development

#### Training Data and Model Architecture

To enable automated detection of waterbirds in drone imagery, we trained a YOLOv10 model (Wang et al. 2024) using Chester Island imagery originally collected in 2021 and reported in Kabra et al. (2022). All model training and evaluation were conducted using the Ultralytics code repository (https://github.com/ultralytics/ultralytics).

Although the 2021 imagery was previously annotated for waterbird detection, we annotated that imagery again to conform to a revised classification scheme of 28 waterbird classes (Table 1). Specifically, we aimed to resolve individual tern species that were previously grouped into a single “Mixed Tern” class, which included *Thalasseus maximus* (Royal Tern; ROYT), *T. sandvicensis* (Sandwich Tern; SATE), *Hydroprogne caspia* (Caspian Tern; CATE), and *Sterna forsteri* (Forster’s Tern; FOTE). The re-annotation closely followed the computer-assisted workflow described in the Annotation Workflows section, except that the original Kabra et al. (2022) annotations served as pre-annotations that annotators reviewed and corrected.

**Table 1.** Classification scheme for the YOLOv10 object detection model. The 28 original annotation classes from the Kabra et al. (2022) Chester Island dataset were consolidated into 15 model classes to address class imbalance in training data. Species are listed in phylogenetic order following the American Ornithological Society Checklist of North and Middle American Birds (Chesser et al. 2024). Non-taxonomic categories (Flying, Egg) appear at the end. Consolidation rationale: (a) annotation sample size too small to support a separate class, so combined with a related species or group; (b) uncommon or non-breeding classes not targeted for species-level detection but annotated during training to reduce model confusion with focal species; (c) white-plumaged wading birds across Ardeidae and Threskiornithidae grouped because visual similarity in nadir drone imagery prevented reliable species-level identification, even for expert annotators; (d) non-bird class retained in training data but not a target for detection in this study.

| Species/Group | Original Annotation Class<br>(n = 28) | Model Class<br>(n = 15) | Consolidation Rationale |
| --- | --- | --- | --- |
| *Leucophaeus atricilla* (Laughing Gull) — Breeding | Laughing Gull — Breeding | LAGU | — |
| *Leucophaeus atricilla* (Laughing Gull) — Nonbreeding | Laughing Gull — Nonbreeding | LAGU | a |
| *Hydroprogne caspia* (Caspian Tern) | Caspian Tern | CATE | — |
| *Sterna forsteri* (Forster's Tern) | Forster's Tern | Tern Spp. | a |
| *Thalasseus maximus* (Royal Tern) | Royal Tern | ROYT | — |
| *Thalasseus sandvicensis* (Sandwich Tern) | Sandwich Tern | SATE | — |
| Terns not identifiable to species | Tern Species | Tern Spp. | — |
| *Rynchops niger* (Black Skimmer) | Black Skimmer | Other | b |
| Shorebird species | Shorebird Species | Other | b |
| Cormorant species | Cormorant Species | Other | b |
| *Pelecanus occidentalis* (Brown Pelican) — Adult | Brown Pelican — Adult | BRPE-Adult | — |
| *Pelecanus occidentalis* (Brown Pelican) — Chick | Brown Pelican — Chick | BRPE-Chick | — |
| *Pelecanus occidentalis* (Brown Pelican) — Nonbreeding | Brown Pelican — Nonbreeding | BRPE-Adult | a |
| *Egretta tricolor* (Tricolored Heron) | Tricolored Heron | TRHE | — |
| *Egretta rufescens* (Reddish Egret) | Reddish Egret | REEG | — |
| *Egretta thula* (Snowy Egret) | Snowy Egret | White Wader | c |
| *Nyctanassa violacea* (Yellow-crowned Night Heron) | Yellow-crowned Night Heron | Other Spp. | b |
| *Nycticorax nycticorax* (Black-crowned Night Heron) | Black-crowned Night Heron | BCNH | — |
| *Ardea alba* (Great Egret) | Great Egret | White Wader | c |
| *Ardea ibis* (Western Cattle-Egret) | Cattle Egret | White Wader | c |
| *Ardea herodias* (Great Blue Heron) | Great Blue Heron | GBHE | — |
| Dark wading birds not identifiable to species | Dark Wader | Other Spp. | b |
| *Eudocimus albus* (White Ibis) | White Ibis | White Wader | c |
| *Platalea ajaja* (Roseate Spoonbill) | Roseate Spoonbill | ROSP | — |
| White-plumaged wading birds not identifiable to species | White Wader | White Wader | c |
| Passerine species | Passerine Species | Other | b |
| Any bird in flight | Flying | Flying | — |
| Visible eggs | Egg | Other | d |

The final revised dataset consisted of 11,500 waterbird annotations, with a long-tailed class distribution dominated by two classes: Royal Tern (4,475 birds; 39%) and Laughing Gull (2,525 birds; 22%) (Figure 3). Given the challenges of training object detectors on imbalanced datasets (Li et al. 2020, Yu et al. 2021), underrepresented species were consolidated into three hierarchical categories: “Tern Species” (terns unidentifiable to species and *Sterna forsteri* [Forster’s Tern], <20 samples); “White Wader” (white wading birds including *Ardea alba* [Great Egret], *Egretta thula* [Snowy Egret], *Ardea ibis* [Western Cattle-Egret], and *Eudocimus albus* [White Ibis]); and “Other Species” (rare taxa with <50 samples including cormorants and shorebirds). The final classification schema used for object detection model development consisted of 15 waterbird classes (Table 1).

**Figure 3.**
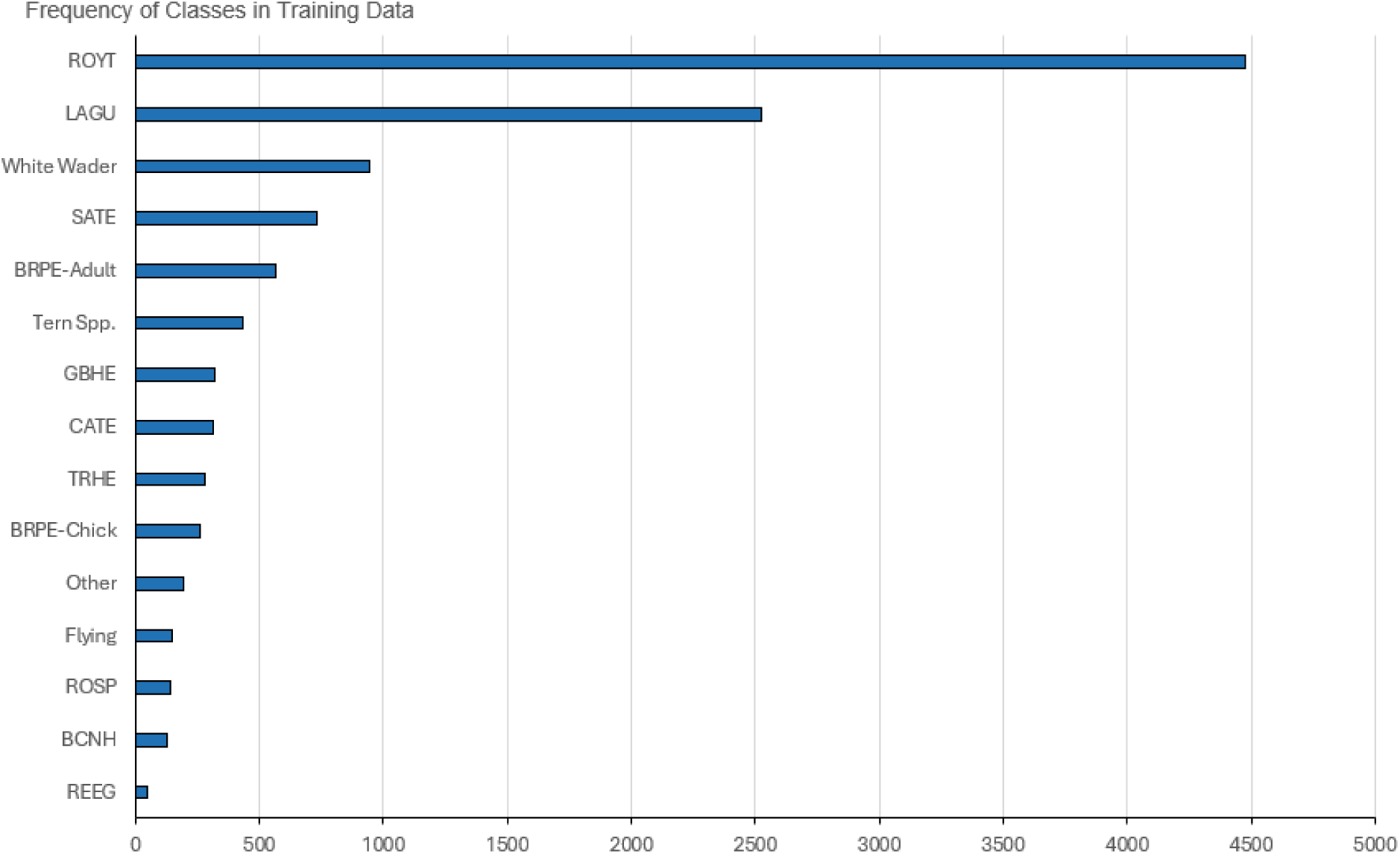
Distribution of 11,500 waterbird annotations across species classes in the 2021 Chester Island training dataset after consolidation from 28 original annotation classes. Two classes, *Thalasseus maximus* (ROYT) and *Leucophaeus atricilla* (LAGU), comprised 61% of all annotations. Classes with fewer than 50 annotations were consolidated into broader categories (Table 1) to address training data imbalance.

*Alt Text: Horizontal bar chart of annotation counts across 15 waterbird classes, sorted from most to least frequent. Royal Tern and Laughing Gull are by far the largest bars; the remaining classes decrease sharply*.

To train and validate the model, the dataset was split 80/20 at the image level to avoid data leakage. Model training was conducted for 20 epochs using the AdamW optimizer (Loshchilov and Hutter 2019) with a learning rate of 0.000526 and momentum of 0.9 (Ultralytics default parameters). The trained model is publicly available. We report validation set performance using the interpolated average precision (AP) metric at an Intersection-over-Union (IoU) threshold of 0.5 (AP@IoU=0.5), following the PASCAL VOC evaluation protocol (Everingham et al. 2010).

### Image Tiling and Pre-/Post-Processing

For model training, the rendered orthomosaic tiles of 10,000 x 10,000 pixels are far too large to process directly. Therefore, these tiles were cropped into 640 x 640-pixel windows using a sliding-window approach with a stride of 350 pixels, following standard practice for object detection in high-resolution aerial imagery. For model evaluation, to avoid double-counting birds across overlapping tiles, all annotations were reprojected back to the original 10,000 x 10,000-pixel tile coordinate space, and non-maximum suppression (NMS) was applied with an IoU threshold of 0.5 (Hosang et al. 2017).

### Drone Imagery Annotation Workflows

All annotation workflows were conducted within LabelStudio (HumanSignal, San Francisco, California, USA), a web-based data labeling platform that supports both manual annotation and ML model integration for pre-labeling. Prior to annotation, the 10,000 × 10,000-pixel orthomosaic tiles were further cropped into 800 × 800-pixel tiles using a sliding-window approach with a stride of 600 pixels (Figure 4). Tiling was necessary for the Human+ML and ML-only workflows because the YOLOv10 architecture cannot process 10,000 × 10,000-pixel tiles directly. The Manual workflow was also tiled at this resolution so that all three workflows operated on identical inputs, enabling direct method comparison. Model outputs from the 800 × 800-pixel tiles could be used directly without requiring reprojection back to the larger tile coordinate space. Although this tile size and stride differed from those used during model training (640 × 640 pixels, stride 350), we found that this sped processing up and had minimal impact on model detection.

**Figure 4.**
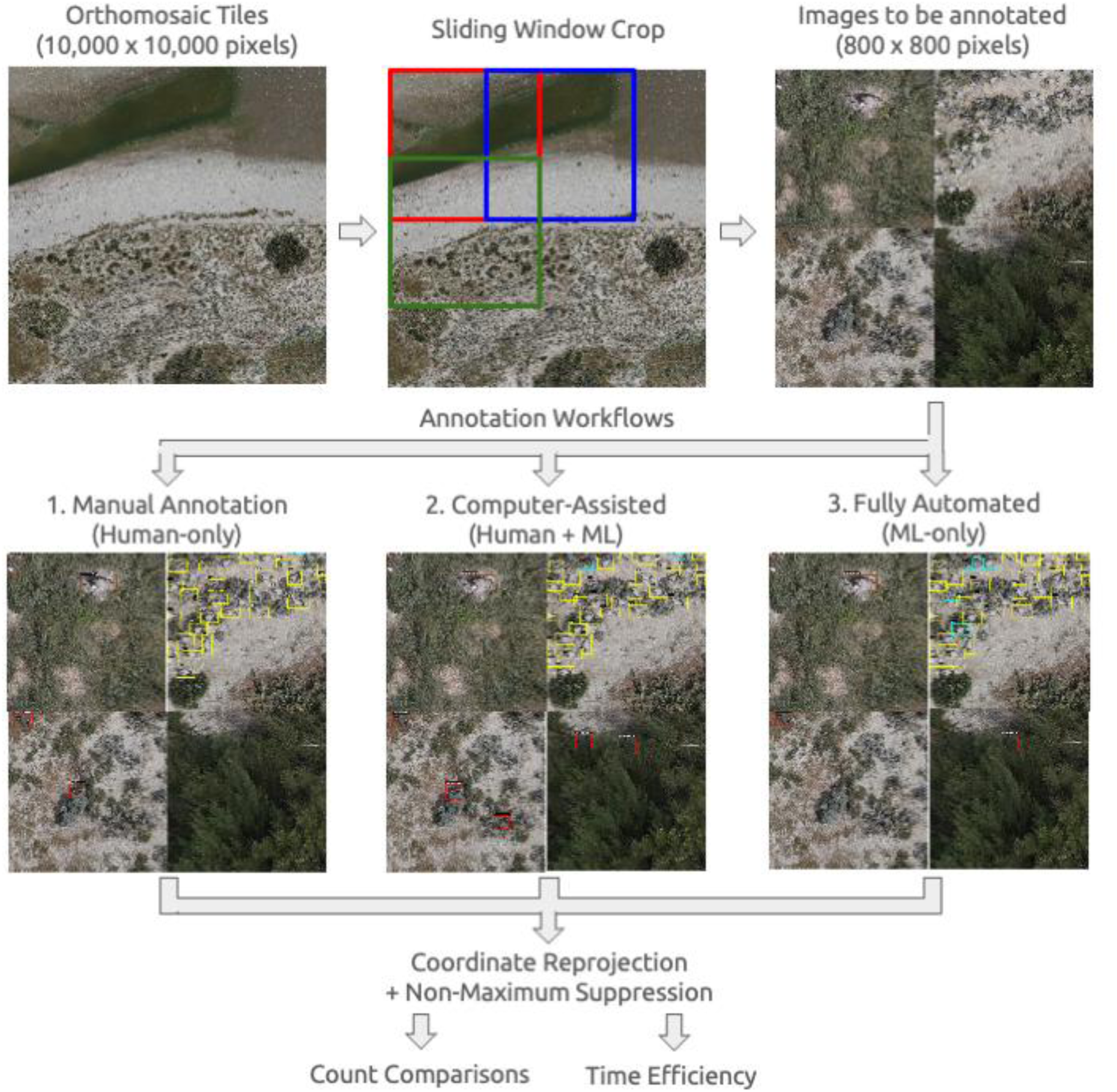
Image processing and annotation workflow. Drone-derived orthomosaic tiles (10,000 × 10,000 pixels) were divided into overlapping 800 × 800-pixel tiles using a sliding-window crop to ensure that birds spanning tile edges were captured in at least one adjacent tile. Tiles were then processed through three parallel annotation workflows: (1) Manual Annotation, in which human annotators drew bounding boxes and assigned species identifications without model assistance; (2) Computer-Assisted (Human + ML) annotation, in which a YOLOv10 object detection model pre-labeled birds and humans verified, corrected, and supplemented detections; and (3) Fully Automated (ML-only) annotation, in which the model produced all bounding boxes and species predictions without human review. Outputs from all three workflows were reprojected to original orthomosaic coordinates and passed through Non-Maximum Suppression to remove duplicate detections in tile-overlap regions, yielding final bird counts and per-method time estimates used in downstream method comparisons.

*Alt Text: Flowchart of the image processing pipeline. A large orthomosaic tile is divided into smaller overlapping tiles, each 800 x 800 pixels, which then branch into three parallel annotation workflows: manual annotation, computer-assisted, and fully automated. These three workflows then merge into a final post-processing step that produces bird counts and timing estimates*.

### Manual Annotation

In the manual workflow, annotators reviewed each 800 x 800-pixel tile independently and performed both localization and classification from scratch. For each visible bird, annotators drew a bounding box around the individual and assigned it to one of the 15 waterbird classes (Table 1). No model-generated pre-annotations were provided. This workflow served as the primary reference standard for evaluating the two automated approaches.

### Computer-Assisted (Human+ML) Annotation

In the computer-assisted workflow, the trained YOLOv10 model was first applied to each tile to generate pre-annotations, which were then loaded into LabelStudio for human review. Annotators were asked to: (1) delete incorrect bounding boxes that did not correspond to a bird, (2) correct misclassified species labels, and (3) add bounding boxes for any birds missed by the model. This workflow decouples localization and classification, allowing annotators to focus their effort on verification and correction rather than on annotation from scratch. Tiles in which the model detected no birds were not forwarded for human review, which substantially reduced the number of tiles requiring human attention.

### Fully Automated (ML-only) Annotation

In the fully automated workflow, the trained YOLOv10 model was applied to all tiles without any human review or correction. Model outputs, i.e., bounding boxes and species classifications, were used directly as the final annotations. No annotator time was required beyond the initial model inference. ML-only processing time was recorded as the total computational time for tiling and model inference, and does not include the time required to develop and train the underlying model.

## Data Analysis

### Count Comparisons

The primary metrics for evaluating the monitoring and annotation approaches are the final counts per species in the surveyed area, which ultimately constitute a major output of monitoring efforts used for wildlife management policies. To collect counts per species in the surveyed area from the drone imagery annotations, we followed the same post-processing protocol used for object detection model evaluation: all bounding boxes were reprojected into a common coordinate space, and NMS was applied with an IoU threshold of 0.5 to remove duplicate detections arising from tile overlap. The final result was a bounding box and class for each bird observed in the original orthomosaic tile, which we then tallied per class across all tiles to obtain the final counts per species in the surveyed area.

### Time Efficiency

A secondary metric for evaluation is the time efficiency, which captures the practical burden of each workflow on practitioners and is directly relevant to the scalability of each approach for operational monitoring programs. Annotation time for the manual and Human+ML workflows was self-reported by annotators, who recorded their active working time in LabelStudio at the conclusion of each session, excluding time spent logging in to the platform or taking breaks. For the ML-only workflow, processing time was recorded as the total wall-clock time for tiling and model inference, and does not include the upfront investment of model development and training.

### Confusion Matrix Analysis

Because drone-based annotations produce pixel-referenced bounding boxes, individual detections from the Human+ML and ML-only workflows can be spatially matched to manual annotations, enabling evaluation of classification accuracy at the level of individual birds rather than aggregate counts. For each drone-based automated method, we matched model-generated bounding boxes to manual annotations using an IoU threshold of 0.5. Matched detections were classified as correct if the assigned species label agreed with the manual annotation, or as a misclassification if labels differed. Unmatched manual annotations were recorded as missed detections (false negatives), and unmatched model detections were recorded as false positives. Results are summarized as a confusion matrix for each method, reporting the number of individuals correctly classified, misclassified by species, and missed entirely. This per-detection analysis complements the count comparisons above, as aggregate count similarity can obscure offsetting errors in detection and classification.

## RESULTS

### Training Data Annotation Effort and Validation

The training data annotation effort for the 2021 Chester Island imagery required approximately 13 hr to annotate 11,500 waterbirds across 62 rendered orthomosaic tiles. Table 2 contains the per-class and overall performance of the trained model on the validation set, measured by AP@IoU=0.5. The model achieves an overall performance of 0.748, with all but 4 classes achieving AP scores above 0.800. Supplementary Figure S1 indicates that Caspian Tern and Tern Spp. are often misclassified as Royal Tern, Reddish Egret are most often misclassified as Other Spp., and many Other Spp. are completely missed (detected as background).

**Table 2:**
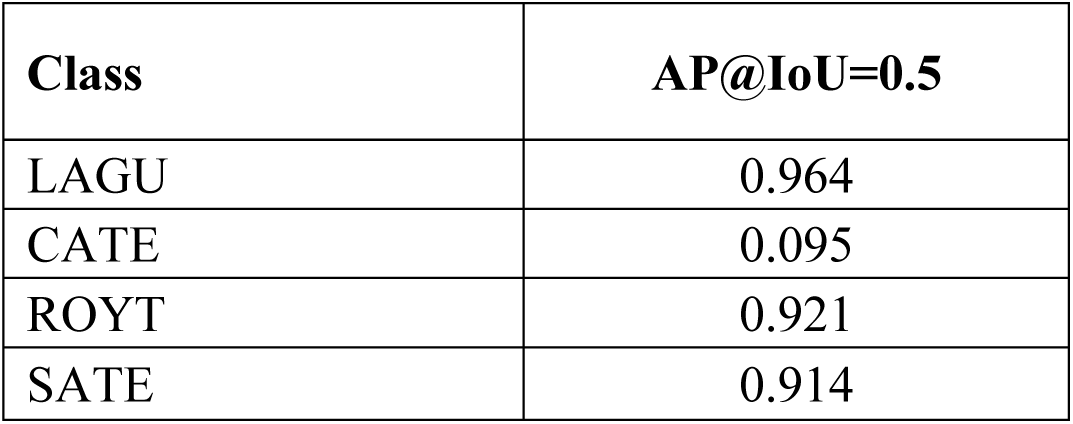
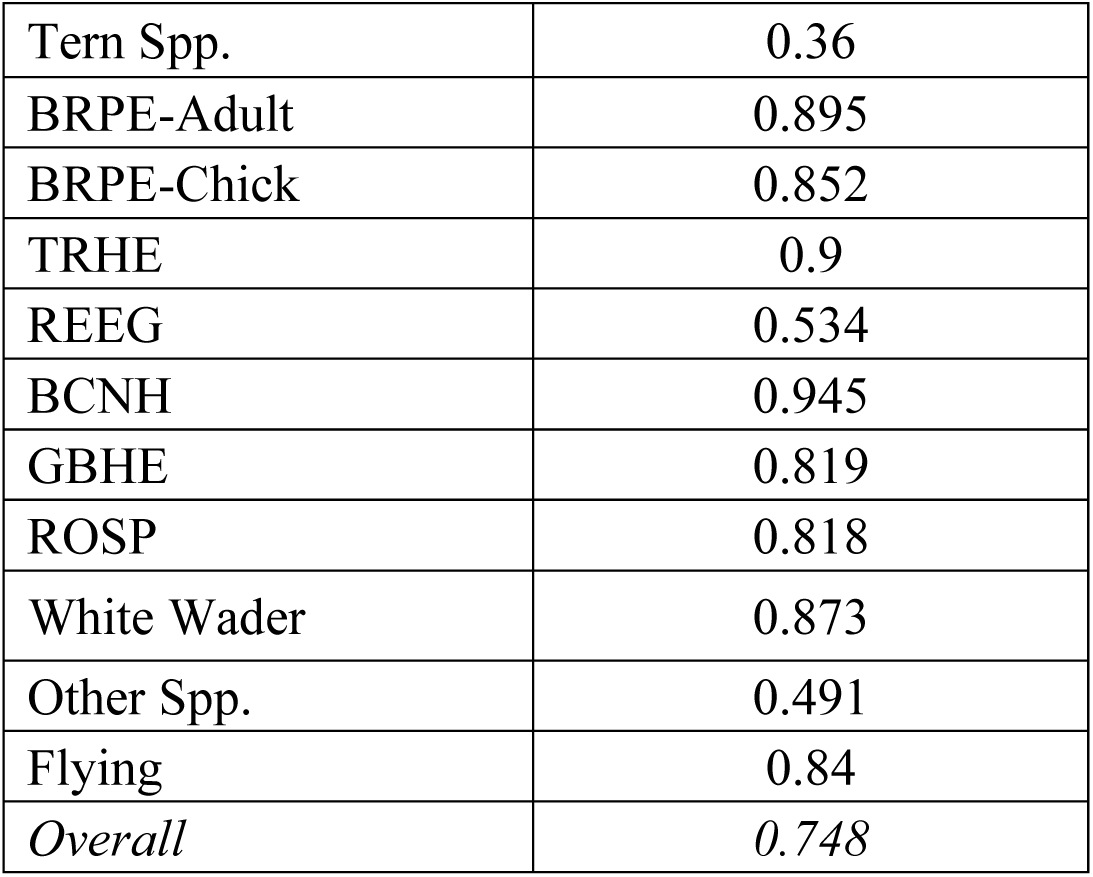
Per-class validation performance of the YOLOv10 object detection model on the 20% held-out 2021 Chester Island validation set, measured as average precision at IoU threshold 0.5 (AP@IoU=0.5).

| Class | $AP@IoU=0.5$ |
| --- | --- |
| LAGU | 0.964 |
| CATE | 0.095 |
| ROYT | 0.921 |
| SATE | 0.914 |
| Tern Spp. | 0.36 |
| BRPE-Adult | 0.895 |
| BRPE-Chick | 0.852 |
| TRHE | 0.9 |
| REEG | 0.534 |
| BCNH | 0.945 |
| GBHE | 0.819 |
| ROSP | 0.818 |
| White Wader | 0.873 |
| Other Spp. | 0.491 |
| Flying | 0.84 |
| *Overall* | *0.748* |

### Count Comparisons Across Methods

Overall detection counts varied across the four monitoring methods. Manual drone annotation, which served as the primary reference standard for drone-based comparisons, detected 6,530 individual birds across the two study polygons. Human+ML detected 5,826 birds (89% of manual), ML-only detected 5,679 birds (87% of manual), and ground counts recorded 5,868 birds for comparable species (90% of manual). Although overall totals were broadly comparable across methods, species-level patterns revealed substantial variation in detection counts.

*Alt Text: Grouped bar charts of bird counts per species across four monitoring methods. Panel A shows ground-nesting species; Panel B shows tree-nesting species along with Other and Flying classes*.

Species-level counts differed considerably across methods, with patterns varying by species (Figure 5). For ground-nesting colonial species, ML-only detection was closest to manual counts for *Thalasseus maximus* (Royal Tern; 98% of manual) but substantially lower for *Leucophaeus atricilla* (Laughing Gull; 72%) and *Pelecanus occidentalis* (Brown Pelican) adults (81%) and chicks (25%). *Thalasseus sandvicensis* (Sandwich Tern) was overdetected by ML-only at 124% of manual. Ground counts exceeded manual drone counts for *T. maximus* (2,675 vs. 1,758) but fell below manual counts for *T. sandvicensis* (1,125 vs. 1,500) and recorded zero Brown Pelican chicks compared to 418 in manual drone annotations. For wading birds, manual drone annotation detected nearly twice as many White Wader group individuals as ground counts (445 vs. 232), while ground counts substantially exceeded manual drone detections for *Egretta tricolor* (Tricolored Heron; 375 vs. 290), *Ardea herodias* (Great Blue Heron; 12 vs. 1), and *Nycticorax nycticorax* (Black-crowned Night Heron; 20 vs. 4).

**Figure 5.**
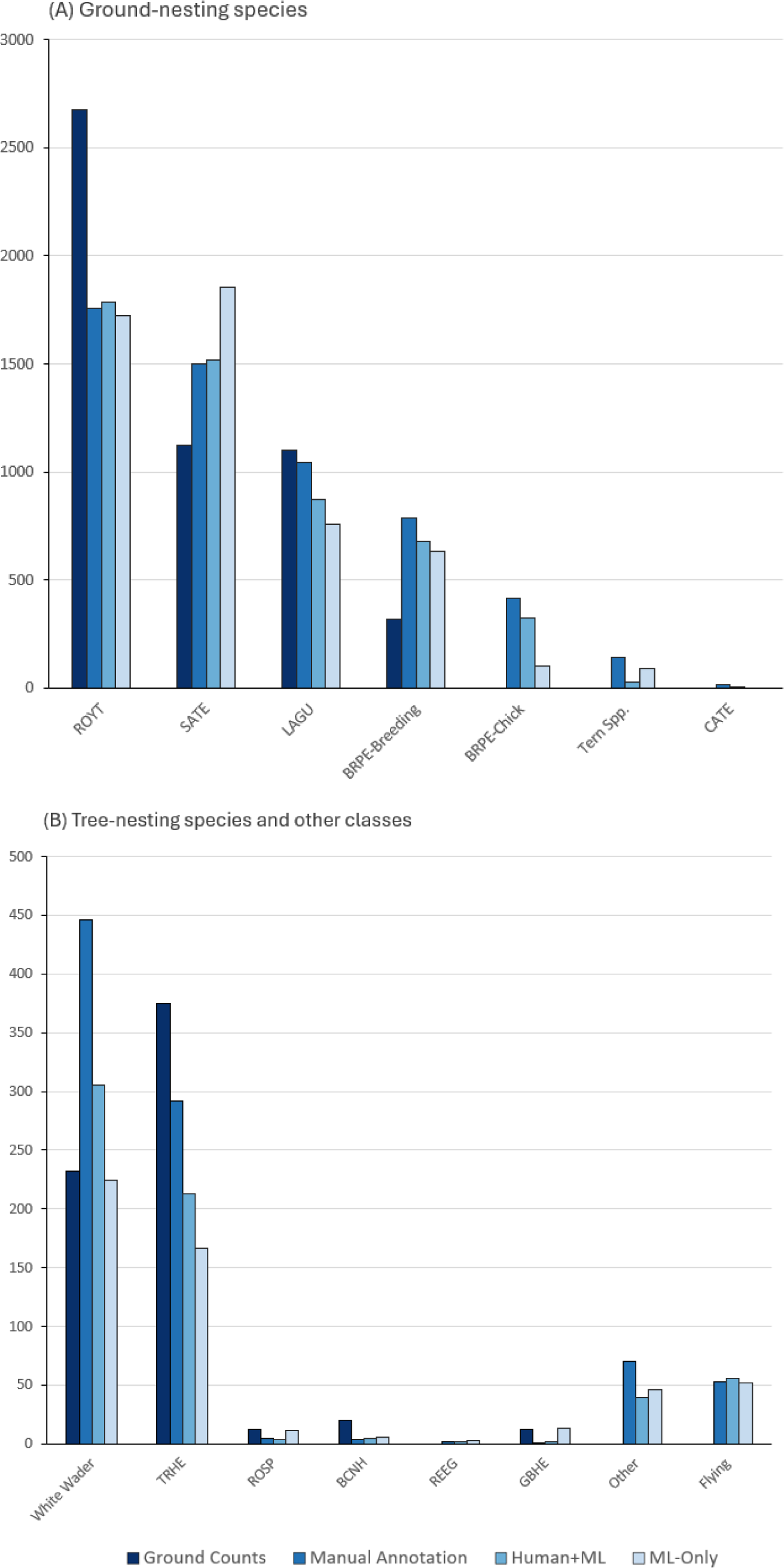
Species-level count comparisons across four waterbird monitoring methods at Chester Island, Texas, 2025: Manual Annotation (drone imagery, reference standard for drone-based comparisons), ML-only (fully automated detection), Human+ML (model output with human verification), and Ground Counts (field-based census). Species are grouped by abundance to allow visualization of both common and less abundant taxa. (**A**) Ground-nesting species, including the most observed species in the imagery. (**B**) Ground-nesting species, the less frequently seen species in the imagery, along with the “Other” and “Flying” classes. See Table 1 for species code definitions and classification scheme.

Ground and manually annotated drone methods recorded notably different tern totals. Ground observers estimated 3,800 terns in total (2,675 *T. maximus* and 1,125 *T. sandvicensis*) and did not use a “Tern Spp.” category. Manual drone annotation identified 1,758 Royal Terns, 1,500 Sandwich Terns, 15 *Hydroprogne caspia* (Caspian Tern), and 140 individuals classified as “Tern Spp.,” for a combined drone total of 3,413 terns, 11% fewer than ground counts.

Unlike the count-based comparisons above, which compare aggregate totals per method, the confusion matrix evaluates classification accuracy on a per-detection basis by matching individual AI detections to manual annotations (Figure 6). Of 6,530 manually annotated individuals, the ML-only model correctly detected and classified 4,443 (68%). The model’s primary source of error was failing to detect birds entirely, with 1,817 individuals (28%) absent from ML-only detections. Misclassification among detected birds was comparatively rare, affecting 270 individuals (4% of manual annotations). The most frequent misclassification was *Thalasseus maximus* classified as *T. sandvicensis* (*n*=78), while the reverse error was uncommon (*n*=6), indicating that confusion between these species was largely unidirectional. *Pelecanus occidentalis* chicks showed secondary misclassification as adult Brown Pelicans (*n*=10) and White Wader group individuals (*n*=21).

**Figure 6.**
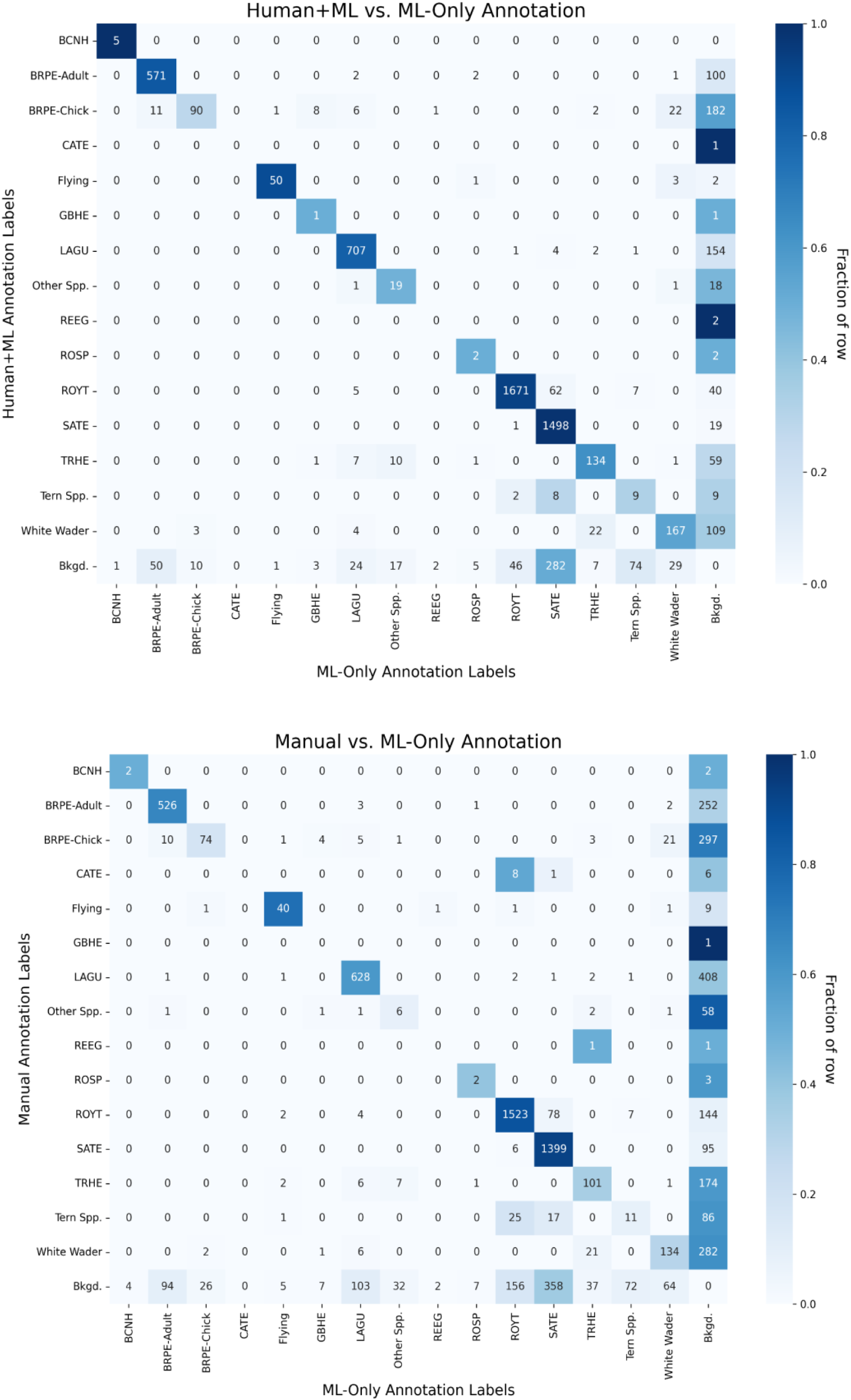
Confusion matrices comparing per-detection classification accuracy across drone-based annotation methods for Chester Island, Texas, 2025. (**A**) Human+ML detections compared against ML-only detections, illustrating species-level corrections introduced by human review. (**B**) Manual annotations (reference standard) compared against ML-only detections. The “Bkgd.” row indicates annotations not matched to an ML-only detection (false negatives); the “Bkgd.” column indicates ML-only detections with no corresponding match in the reference workflow (false positives). See Table 1 for species code definitions.

Human review of ML detections increased both total counts and classification accuracy. Human+ML detected 5,826 individuals compared to 5,679 for ML-only, recovering an additional 147 detections. Species-level gains were greatest for *Pelecanus occidentalis* chicks (103 ML-only vs. 323 Human+ML) and White Wader group individuals (224 vs. 305). Human review also reduced the Sandwich Tern overcount from 1,854 (ML-only) to 1,518 (Human+ML), more closely aligning with the manual annotation total of 1,500. Despite these improvements, Human+ML counts remained below manual annotation totals for most species.

*Alt Text: Two confusion matrices. Diagonal cells demonstrate the detections that were correctly classified. The darker the cell, the more overlap between the two methods. Panel A compares Human plus Machine Learning detections to Machine Learning only; Panel B compares Manual annotations to Machine Learning only*.

### Time Efficiency

Annotation time differed substantially across drone-based methods (Table 3). Manual drone annotation of the two polygons (37 10k x 10k-pixel orthomosaic tiles, 10,693 800 x 800-pixel tiles) required 2,429 min (40.5 hr), or about 161 birds per hour. Human+ML annotation of the same image set required 463 min (7.7 hr), an 81% reduction in annotation time or about 755 birds per hour. ML-only processing required approximately 46 min across the same 37 images (64 s for tiling and 11 s for model inference per image on a free-tier GPU), a 98% reduction relative to manual annotation or about 7,367 birds per hour. These times reflect active annotation only and exclude time spent logging into the annotation platform or annotator breaks. ML-only times exclude the computational resources and expertise required to develop and train the underlying detection and classification models.

**Table 3.** Annotation time, detection counts, and percent reduction relative to manual annotation for three drone-based workflows applied to two survey polygons on Chester Island, Texas, 2025. Manual annotation served as the reference standard for drone-based comparisons. ML-only processing was conducted on a free-tier Google Colab T4 GPU (Google, Mountain View, California, USA) and excludes upfront model training time.

| Method | Total time | Birds detected | % of manual count | Time reduction |
| --- | --- | --- | --- | --- |
| Manual | 40.48 hr | 6,530 | 100% | — |
| Human+ML | 7.72 hr | 5,826 | 89.20% | 80.90% |
| ML-only | ~46 min | 5,679 | 87.00% | 98.10% |

## DISCUSSION

This study represents one of the first systematic comparisons of manual, semi-automated, and fully automated approaches for counting waterbirds in drone imagery from a large, mixed-species colony. Semi-automated annotation, combining automated detection with human classification, captured 89% of manually identified waterbirds while requiring only 19% of the annotation time. Fully automated processing detected 87% of birds in approximately 46 min of computation. These results demonstrate that CNN-based detection and classification can be applied to a complex, mixed-species colony supporting at least 15 waterbird classes, including visually similar tern species that challenge both human annotators and machine learning models. Previous automated counting efforts have primarily targeted single-species colonies or assemblages with few visually distinct species (Chabot and Francis 2016, Hong et al. 2019, Hayes et al. 2021). By comparing all four approaches on the same survey areas during the same breeding season, including concurrent ground surveys used by the Texas Colonial Waterbird Survey for over 50 years (Ortego et al. 2011), we were able to isolate the contributions of automated detection from automated classification. This distinction has direct practical implications: the semi-automated workflow effectively simulates a generalized bird detector that locates individuals regardless of species, representing a practical entry point for programs seeking efficiency gains without the investment required to train species-level classifiers.

### Data

The YOLOv10 model applied in this study was trained on 2021 Chester Island annotations that originally contained 28 species, consolidated to 15 classes to address class imbalance in the training data (Kabra et al. 2022). This consolidation grouped underrepresented species into broader categories such as “White Wader”, and “Other Spp.”, which improved model training stability but limited the taxonomic resolution of fully automated classifications for those species. The training dataset was heavily skewed toward the colony’s most common species, with Royal Tern and Laughing Gull comprising the majority of annotations, while several classes contained fewer than 50 training samples. This imbalance has direct implications for species-specific model performance discussed below.

For this project, manual annotation served as the reference standard for evaluating both Human+ML and ML-only, but this benchmark is itself imperfect. Our annotator comparison analysis found that detection agreement among three independent annotators was high (99%), suggesting that trained observers reliably locate birds in drone imagery. Classification agreement, by contrast, averaged only 78% across annotator pairs, with most disagreement concentrated among morphologically similar tern species and reflecting differences in hierarchical taxonomic specificity rather than outright misidentification (Supplementary Figure S2). The detection rates reported below for Human+ML and ML-only should therefore be interpreted as performance relative to a consistent and relatively complete, but imperfect, benchmark. This is a common challenge in ecological monitoring, where the assumed source of truth has historically been data from field observations. Even so, manually annotated drone imagery represents a substantial improvement over traditional survey methods for ground-nesting species. Ground-based surveys have been shown to produce less precise count estimates than aerial surveys (Hodgson et al. 2016, 2018; Prosser et al. 2023), and boat-based surveys risk missing nests due to low vantage points (Wetlands International 2010). Preliminary comparisons of boat-based counts and manually annotated drone imagery at Evia Island in Galveston Bay, Texas, which motivated the present study, illustrate this gap: boat crews counted 705 Brown Pelican compared to 928 in drone imagery (76%), 1,950 Laughing Gull compared to 11,535 (17%), and 2,655 terns compared to 8,773 (30%) (authors’ personal observation).

### Model Performance

Overall detection rates were broadly comparable across the four monitoring methods, with Human+ML detecting 89% of manually annotated birds, ML-only detecting 87%, and ground counts recording 90% of all species. However, these similar totals mask substantial species-level variation driven by nesting ecology, morphology, and training data representation. For ground-nesting colonial species, including the four most common species (Royal Tern, Sandwich Tern, Brown Pelican, and Laughing Gull) where the model had abundant training data, ML-only detection ranged from 72% (Laughing Gull) to 124% (Sandwich Tern) of manual counts, with Royal Tern at 98% and Brown Pelican adults at 81%. For wading birds that nest in or beneath vegetation, the most reliable method changes. Ground counts substantially exceeded drone-based detections for Great Blue Heron (12 vs. 1 in manual drone annotation), Black-crowned Night Heron (20 vs. 4), and Tricolored Heron (375 vs. 290), while manual drone annotation detected nearly twice as many White Wader group individuals as ground counts (445 vs. 232). These patterns suggest that no single method provides a complete census across all waterbird species that nest along the Texas coast.

Two types of comparisons are used throughout this section. Count-based comparisons evaluate aggregate species totals across methods, where over-detections and missed detections can offset each other, producing similar overall totals even when individual-level accuracy varies. The confusion matrix, by contrast, tracks what happened to each individual bird by matching ML-only detections to manual annotations on a per-detection basis. This is why, for example, Royal Tern can appear at 98% of manual counts while the overall per-detection correct classification rate across all species is 68% (4,443 of 6,530).

The model performed well on the colony’s two dominant tern species despite their similar appearance in drone imagery (Figure 7). Royal Tern was detected at 98% of manual counts, and Sandwich Tern at 124%. Of 1,758 manually annotated Royal Terns, only 78 (4%) were misclassified as Sandwich Tern in the ML-only confusion matrix, and the reverse error was even rarer (6 of 1,500 Sandwich Terns). This strong performance likely reflects the abundance of both species in the training data, with Royal Tern alone comprising 43% of all training annotations. The Sandwich Tern overcount suggests that the model occasionally assigned other tern species to Sandwich Tern, but human review in the Human+ML workflow corrected most of this, reducing the count from 1,854 to 1,518 and closely aligning with the manual total of 1,500. Kabra et al. (2022) achieved strong detection performance for terns but grouped all tern species into a single “Mixed Tern” class, explicitly identifying species-level separation as a key challenge for future work. Arnberg et al. (2026) similarly grouped terns into a single class and reported an F1 score of 0.97. That our model achieved strong performance while classifying Royal and Sandwich Tern separately represents a direct advance on Kabra et al. (2022) and suggests that species-level tern classification from nadir imagery is feasible when sufficient training data are available for each species. Caspian Tern, however, illustrates the limits of this approach when training data are insufficient. Manual annotation identified 15 Caspian Terns in the study area, but the ML-only model detected none, and even the Human+ML workflow classified only a single individual. The model’s validation performance for this species was effectively zero (*recall* = 0.00, *mAP50* = 0.018), likely reflecting very low training sample size combined with strong morphological overlap with Royal Tern. Our annotator agreement analysis reinforces this: Caspian Tern had the lowest pairwise classification agreement of any species (32%), with one annotator classifying the majority of these individuals as Royal Tern. When even expert human annotators struggle to distinguish Caspian from Royal Terns in aerial imagery, training a model to make this distinction reliably would require substantially more labeled examples or imagery.

**Figure 7.**
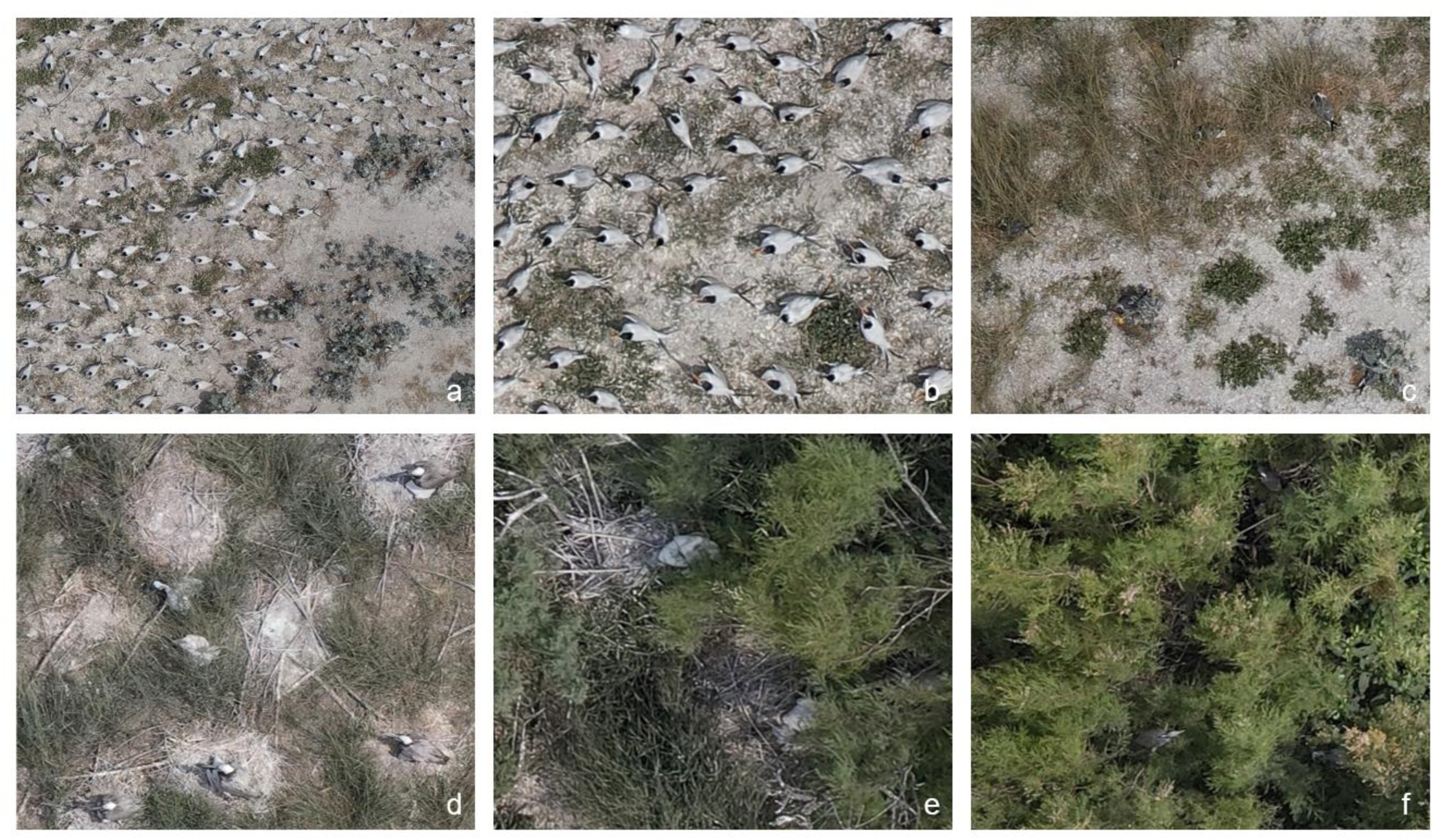
Examples of detection and classification challenges in nadir drone imagery from Chester Island, Texas. (**A**) Dense mixed tern colony on open substrate, where individuals are clearly visible from above. (**B**) *Thalasseus maximus* and *T. sandvicensis* nesting in close proximity, illustrating the morphological similarity. (**C**) *Leucophaeus atricilla* partially obscured by low vegetation, contributing to the 72% detection rate relative to manual annotation. (**D**) *Pelecanus occidentalis* adults and chicks among sparse vegetation, showing the distinct visual profiles of each age class. (**E**) *P. occidentalis* chick appearing as a diffuse white mass on the nest, similar in appearance to white-plumaged wading birds. (**F**) *Egretta tricolor* and a white wader nesting beneath dense canopy, where vegetation occlusion reduced drone-based detection rates.

*Alt Text: Six aerial drone photographs from Chester Island. Panel A: dense mixed tern colony on open ground. Panel B: Royal and Sandwich Terns nesting side by side. Panel C: Laughing Gull nests partly hidden by vegetation. Panel D: Brown Pelican adults and chicks among sparse vegetation. Panel E: Brown Pelican chick appearing as a diffuse white shape on a nest. Panel F: Tricolored Heron and a white wading bird beneath dense canopy*.

Ground and drone methods recorded notably different tern totals. Ground observers estimated 3,800 terns (2,675 Royal Tern and 1,125 Sandwich Tern), while manual drone annotation identified 3,413 terns across all tern classes, 11% fewer. However, the species-level breakdown differed substantially: ground counts recorded nearly 1,000 more Royal Terns than manual drone annotation (2,675 vs. 1,758) but far fewer Sandwich Terns (1,125 vs. 1,500). This discrepancy likely reflects the estimation methods used by ground teams, who estimate total terns and then apply an estimated ratio of Royal to Sandwich Tern rather than counting each species independently. A nearly 1,000-bird overestimate for a single species is considerable, and underscores a key limitation of field estimation methods for densely packed, mixed-species tern groups. Drone annotators, by contrast, classified each individual separately and assigned 140 birds (4% of terns) to “Tern Spp.” when diagnostic features were insufficient for species-level identification, a category not used by ground observers. These differences highlight how estimation methods can introduce species-level biases even when overall totals are broadly similar, and reinforce the value of concurrent surveys using both methods during any transition between monitoring protocols, as recommended by Hodgson et al. (2016).

Brown Pelican performance revealed a different challenge. While the model detected adult Brown Pelicans at 81% of manual counts, it captured only 25% of Brown Pelican chicks, the lowest detection rate of any class. Brown Pelican chicks were underrepresented in the training data, with only 140 annotations compared to 257 for adults. Chicks also present a fundamentally different visual profile than adults, appearing as amorphous white masses in nests rather than the distinctive adult silhouette, often obscured by vegetation or by adults on top of chicks (Figure 7). The confusion matrix confirmed secondary misclassification of chicks as the White Wader group individuals (*n*=21), suggesting that the model struggled to distinguish chick morphology from similarly diffuse light-colored shapes. The Human+ML workflow recovered substantially more chicks (323 vs. 103 for ML-only), suggesting that even imperfect automated detection provides a useful starting point that human annotators can supplement. Note, though, that ground surveys recorded zero chicks in the study area, as approaching pelican nesting zones on foot causes documented adult flushing and chick exposure (Jones et al. 2020) and chicks are, again, often obscured by vegetation or adults.

The comparison between ground-based and drone-based counts for wading birds highlighted complementary strengths tied to nesting ecology. Many wading bird species could not be reliably differentiated in nadir imagery, which is why several were consolidated into the “White Wader” group class for this study (Figure 7). Even at this coarser taxonomic level, manual drone annotation detected nearly twice as many White Wader group individuals as ground counts (445 vs. 232), likely because these species are visible from above when standing at or near nest sites in otherwise dense vegetation. The pattern reversed for species that nest under dense canopy, with ground counts recording substantially more Great Blue Heron (12 vs. 1), Black-crowned Night Heron (20 vs. 4), and Tricolored Heron (375 vs. 290) than drone-based methods. Barr et al. (2019) similarly found that UAS surveys were prone to low detection at sites where vegetation occluded nests. These patterns reinforce that the most reliable count method depends on species ecology, and that no single method provides a complete census across nesting strategies.

### Accuracy vs. Efficiency Trade-Offs

The progression from manual annotation to fully automated processing reveals a clear but non-linear tradeoff between time investment and detection accuracy. Manual annotation required 40.48 hr to process the selected dataset, while Human+ML reduced this to 7.72 hr, an 81% time reduction. ML-only processing completed the effort in approximately 46 min on a free-tier Google Colab T4 GPU (Google, Mountain View, California, USA) (Table 3). These time savings are broadly consistent with those reported elsewhere. Kellenberger et al. (2021) reported that ML-based detection reduced processing time for approximately 21,000 seabirds from three weeks of manual annotation to 4.5 hr. Arnberg et al. (2026) similarly found that deep learning models served as effective preliminary annotators within semi-automated pipelines, enabling experts to validate and correct model outputs rather than annotate from scratch. Our results extend these findings by quantifying the intermediate step: the Human+ML, or human-in-the-loop, workflow, in which a trained model handles detection while a human expert provides species classification. This intermediate approach captured 89% of the birds identified by fully manual annotation while requiring only 19% of the time, representing a practical balance between efficiency and accuracy that has not been explicitly quantified in previous studies.

A fundamental limitation of the Human+ML workflow warrants emphasis. In our tiling workflow, the full-resolution images were divided into 10,693 tiles for annotation (Figure 4). When the bird detector was applied, only tiles in which the model detected at least one bird were forwarded to the human annotator for species classification, reducing the number of tiles requiring human review from 10,693 to 2,020. While this filtering drove the majority of the time savings, it also meant that tiles containing birds the model failed to detect were never reviewed. This tile-level filtering explains the persistent undercount relative to fully manual annotation, in which annotators systematically reviewed all 10,693 tiles regardless of model output. The limitation could be addressed by routing all tiles to human review, but doing so would substantially reduce the time savings that make the semi-automated approach practical. Arnberg et al. (2026), however, noted that the relationship between human and model detection is not strictly one-directional; their early test models identified birds that manual annotators had missed on initial passes, suggesting that human and automated approaches have complementary detection strengths that a well-designed workflow could exploit.

### Considerations

Several broader considerations should inform how practitioners evaluate these tools for their own monitoring programs. First, our model was trained and applied at sites exclusively along the coast of Texas, with many of the same species, consistent flight protocols, and similar habitats. Performance on other colonies with different species assemblages, sensor types, vegetation structure, or substrates is unknown. Even colonies along the Texas coast that have different dominant species or species that aren’t common in our training data may have different classification results and may require retraining or, at a minimum, revalidation before these results can be expected. However, a consistent finding across our results is that bird detection was substantially more generalizable than species classification. The bird detector performed well across all 15 classes regardless of training data representation, and the Human+ML workflow captured 89% of manually annotated birds while requiring only 19% of the time. This pattern aligns with Weinstein et al. (2022), who showed that general bird detection models can perform well in novel environments without local training data. This suggests that detection models may generalize across sites more readily than classification models.

The ML-only processing time of approximately 46 min for 37 images reflects a specific hardware configuration (free-tier Google Colab T4 GPU) will vary with available computational resources. Practitioners with access to more powerful hardware would see faster processing, while those relying on CPU-only systems would see substantially slower results. Additionally, the efficiency comparisons presented here do not account for the time required to create training data. The 2021 annotations that served as training data for this model required considerable expert effort, and any program adopting a fully automated approach will face this upfront investment before realizing the time savings described above. Weinstein et al. (2022) showed that starting from a general pretrained model reduces both the amount of local training data needed and the computational cost of training, which could substantially lower this barrier for new programs. Finally, our model was trained on 2021 imagery and applied to 2025 survey data, a four-year gap during which colony composition, nesting density, and vegetation conditions may have changed. That the model still performed reasonably well across this gap is encouraging, but ongoing validation against manual annotation will be important to ensure that model performance does not degrade as conditions shift over time.

## Conclusion

This study provides one of the first systematic comparisons of manual, semi-automated, and fully automated approaches for counting waterbirds in drone imagery from a large, mixed-species colony. The four-method comparison framework allowed us to isolate the contributions of automated detection from automated classification, revealing that detection generalizes well across species while classification performance depends heavily on training data availability and morphological distinctiveness. The Human+ML workflow emerged as a practical middle ground, capturing 89% of manually annotated birds while requiring only 19% of the time, and the fully automated pipeline demonstrated strong performance for well-represented species while highlighting clear limitations for rare or visually similar taxa.

Drone-based surveys have been found to reduce disturbance to nesting colonies compared to traditional ground-based methods (Rush et al. 2018, Edney et al. 2023, Leija et al. 2023). Our results demonstrate that this advantage can be paired with processing workflows that are fast enough to make drone surveys practical at scale. The Human+ML workflow, which required less than eight hr to process imagery that took over 40 hr to annotate manually, makes it more feasible to survey sites within a given monitoring window or to allocate biologist time to other conservation priorities. For programs like the Texas Colonial Waterbird Survey, which monitors dozens of islands annually, even modest per-site time savings could allow for adoption of drone-based surveys across these sites and expand monitoring capacity across the region. Standardization is another potential benefit. Traditional ground surveys are variable by nature, dependent on observer experience, vantage point, and estimation methods used. Drone imagery, by contrast, produces a permanent, reviewable record that can be reanalyzed as models improve or as new questions arise. Pairing this archive with consistent automated or semi-automated processing could reduce inter-observer variability and improve comparability of population estimates across sites and years, strengthening the data foundation for conservation decision-making.

The strong performance of automated detection across all species classes points toward a practical path forward: generalized bird detection models that locate individuals regardless of species, paired with either human classification or site-specific classification models as needed. Weinstein et al. (2022) demonstrated that a general bird detection model trained on over 250,000 annotations from 13 diverse projects could detect birds in novel environments without any local training data, and that fine-tuning with as few as 1,000 local annotations substantially improved performance. The progression from fully manual annotation to semi-automated to full automation described in this study provides a practical framework for monitoring programs to adopt AI assistance incrementally, at a pace that matches their resources, expertise, and monitoring objectives.

## Acknowledgments

We would like to thank the Center for Research Computing and the Data to Knowledge Lab at Rice University as well as the many students who helped support this project during their time in the Data to Knowledge Lab, including Kaiyan Ma, Sirui Hao, Akshay Raj, Will Miraglia, and Jason Nguyen. We would also like to thank Marissa Lamb for her support of this project and Gautam Apte for his time annotating images. Finally, we would like to thank HumanSignal for providing Label Studio Enterprise to Rice University as part of their Academic Program.

## Declaration of generative AI in the writing process

The authors used Claude (Anthropic; https://claude.ai) to review writing for clarity and consistency, verify citation accuracy against source materials, and conduct style and grammar checks. The authors reviewed and edited any generated material and accept full responsibility for the content of the publication.

## Ethics statement

All drone surveys of Chester Island were conducted with Audubon Texas research authorization and with FAA Part 107-certified pilots operating in compliance with 14 CFR Part 107. Flight protocols were designed to minimize disturbance to nesting waterbirds, following best-practice guidelines for drone operations near colonial waterbird colonies. No birds were captured, handled, marked, or otherwise physically interacted with during this study; all data were collected via non-invasive aerial imagery. Ground-based surveys conducted as part of the annual Texas Colonial Waterbird Survey followed standardized survey protocols.

## Author contributions

- Conceived the idea, design, experiment (supervised research, formulated question or hypothesis): Anna Vallery, Krish Kabra, Richard Gibbons, Hank Arnold, Arko Barman

- Performed the experiments (collected data, conducted the research): Anna Vallery, Krish Kabra, Richard Gibbons, Hank Arnold, Nicholas Minnich

- Wrote the paper (or substantially edited the paper): Anna Vallery, Krish Kabra, Richard Gibbons, Arko Barman

- Developed or designed methods: Anna Vallery, Krish Kabra, Richard Gibbons, Hank Arnold, Arko Barman

- Analyzed the data: Anna Vallery, Krish Kabra

- Contributed substantial materials, resources, or funding: Hank Arnold

## Data availability

Data used in this study, including 2025 Chester Island imagery, annotation files, and model outputs, will be deposited into the Dryad repository with a citable DOI prior to acceptance. The trained model and weights are currently available on GitHub and will be archived with a DOI prior to acceptance.

