## Supplemental Materials for "Application of Machine Learning Tools for Waterbird Colony Monitoring Provides Gains in Precision and Temporal Efficiency"

**Supplementary Materials**

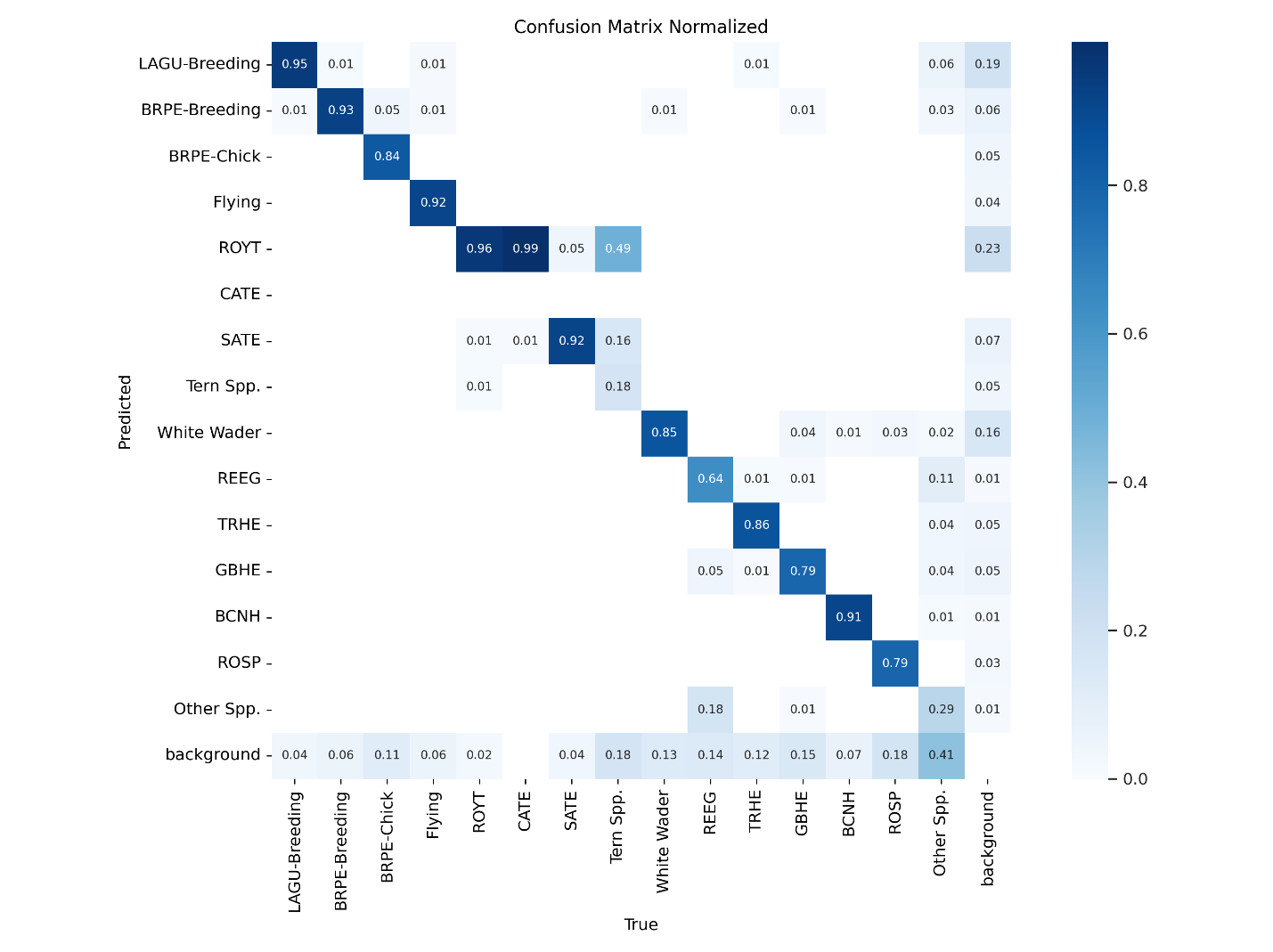

**Supplementary Material Figure S1.** Normalized confusion matrix for the YOLOv10 object detection model evaluated on the 2021 Chester Island validation set (Kabra et al. 2022), with classes consolidated to the 15 model classes described in Table 1. Columns represent true annotation labels and rows represent model predictions; cell values give the proportion of true-class instances assigned to each predicted class (column-normalized), with diagonal values indicating correct classification. The "background" row indicates objects in the ground truth that were not detected by the model (false negatives), and the "Other Spp." class captures uncommon or non-breeding species annotated to reduce confusion with focal species (Table 1). Confusion patterns referenced in the main text are visible here, including misclassification of Caspian Tern (CATE) and Tern Spp. as Royal Tern (ROYT) or Sandwich Tern (SATE), misclassification of Reddish Egret (REEG) as Other Spp., and detection failures for Other Spp.

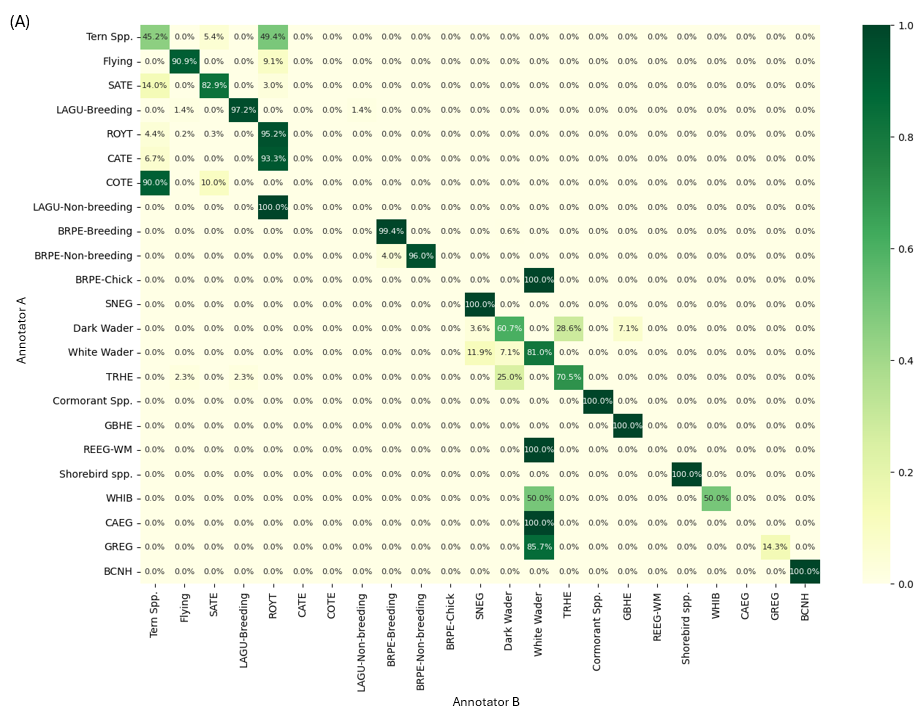

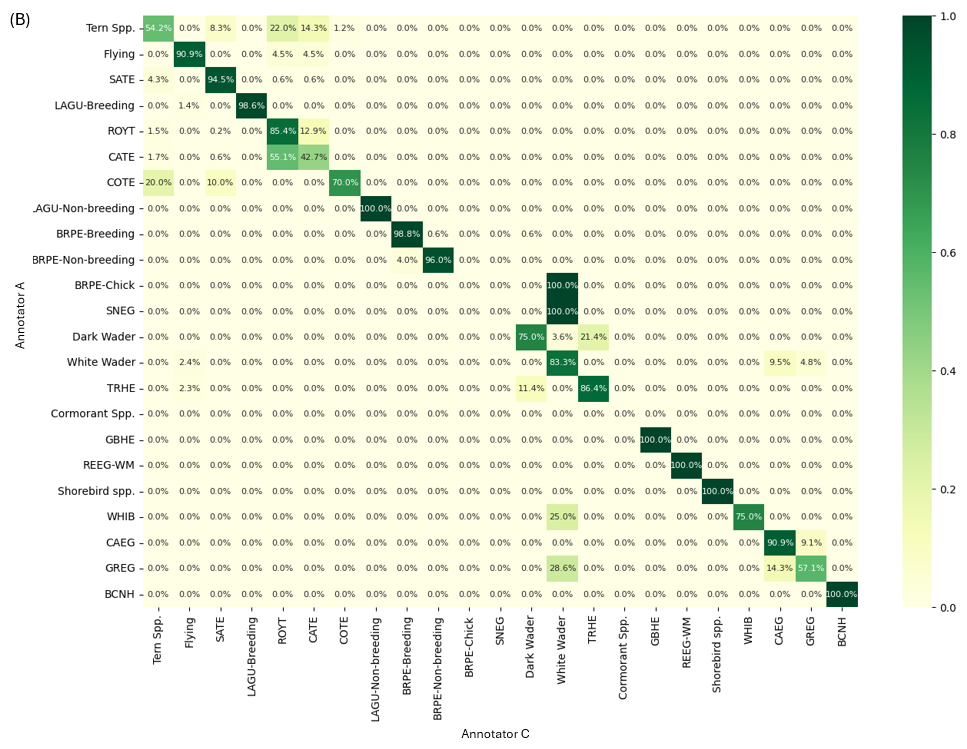

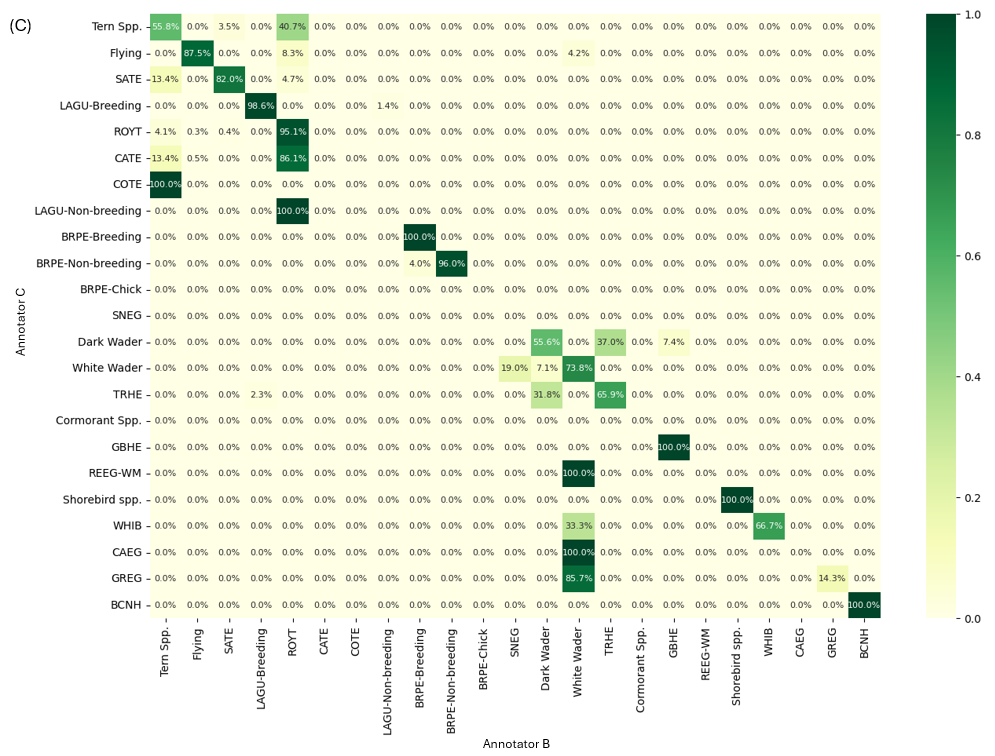

**Supplementary Material Figure S2.** Pairwise classification agreement between three independent annotators of the 2025 Chester Island drone imagery, evaluated at the level of the 28 original annotation classes used during annotation (Table 1) prior to consolidation to the 15 model classes. Each panel shows the agreement between one annotator pair: (A) Annotator A and Annotator B, (B) Annotator A and Annotator C, and (C) Annotator B and Annotator C. Rows and columns represent each annotator's labels for the same subset of images; cell values give the proportion of one annotator's labels assigned to each of the other annotator's classes (row-normalized), with diagonal values indicating agreement. Detection agreement among the three annotators was 99.5%; classification agreement averaged 78% across pairs. Disagreement was concentrated among morphologically similar tern species (Royal Tern, Caspian Tern, Sandwich Tern, and Tern Spp.) and among white-plumaged wading birds (Great Egret, Snowy Egret, Cattle Egret, White Ibis), reflecting differences in hierarchical taxonomic specificity rather than outright misidentification. For example, one annotator assigned a confident species-level identification (e.g. ROYT) when another used a more general group label (Tern Spp.). Many of these disagreements would resolve to matches under the consolidated 15-class scheme used by the model. Abbreviations not defined in Table 1: COTE (*Sterna hirundo,* Common Tern) and REEG-WM (*Egretta rufescens*, white morph Reddish Egret,).
